# Novel enzyme-based reduced representation method for DNA methylation profiling with low inputs

**DOI:** 10.1101/2024.05.24.595803

**Authors:** Qianli Liu, Kathryn A. Helmin, Zachary D. Dortzbach, Carla P. Reyes Flores, Manuel A. Torres Acosta, Jonathan K. Gurkan, Anthony M. Joudi, Nurbek Mambetsariev, Luisa Morales-Nebreda, Mengjia Kang, Luke Rasmussen, Xóchitl G. Pérez-Leonor, Hiam Abdala-Valencia, Benjamin D. Singer

## Abstract

DNA methylation at cytosine-phospho-guanine (CpG) residues is a vital biological process that regulates cell identity and function. Although widely used, bisulfite-based cytosine conversion procedures for DNA methylation sequencing require high temperature and extreme pH, which lead to DNA degradation, especially among unmethylated cytosines. Enzymatic methylation sequencing (EM-seq), an enzyme-based cytosine conversion method, has been proposed as a less biased alternative for methylation profiling. Compared to bisulfite-based methods, EM-seq boasts greater genome coverage with less GC bias and has the potential to cover more CpGs with the same number of reads (i.e., higher signal-to-noise ratio). Reduced representation approaches enrich samples for CpG-rich genomic regions, thereby enhancing throughput and cost effectiveness. We hypothesized that enzyme-based technology could be adapted for reduced representation methylation sequencing to enable high-resolution DNA methylation profiling on low-input samples, including those obtained from clinical specimens. We leveraged the well-established differences in methylation profile between mouse CD4+ T cell populations to compare the performance of a novel reduced representation EM-seq (RREM-seq) procedure against an established reduced representation bisulfite sequencing (RRBS) protocol. While the RRBS method failed to generate reliable DNA libraries when using <2 ng of DNA (equivalent to DNA from around 350 cells), the RREM-seq method successfully generated reliable DNA libraries from 1–25 ng of mouse and human DNA. Ultra-low-input (<2-ng) RREM-seq libraries’ final concentration, regulatory genomic element coverage, and methylation status within lineage-defining Treg cell-specific super-enhancers were comparable to RRBS libraries with more than 10-fold higher DNA input. RREM-seq also successfully detected lineage-defining methylation differences between alveolar Tconv and Treg cells obtained from mechanically ventilated patients with severe SARS-CoV-2 pneumonia. Our RREM-seq method enables single-nucleotide resolution methylation profiling using low-input samples, including from clinical sources.

## Introduction

In mammalian genomes, 60-80% of cytosine-phospho-guanine dinucleotides (CpGs) are modified with a methyl group at the fifth carbon position (5mC).^1^ These CpG dinucleotide residues exhibit a nonuniform distribution throughout the genome and are clustered in GC-rich CpG islands.^2^ In most cases, CpG methylation is associated with transcriptional repression, and CpG islands tend to be hypomethylated in regulatory regions around the transcription start site of actively transcribed genes. Currently, bisulfite-based methods, including whole-genome bisulfite sequencing (WGBS) and reduced representation bisulfite sequencing (RRBS), are considered the gold standard for methylation mapping with single-nucleotide resolution.^3^

Despite its popularity, bisulfite sequencing requires extreme temperature and pH during library preparation, which cause DNA degradation, especially among unmethylated cytosines.^4^ This disproportionate damage of unmethylated cytosines results in libraries with inaccurate GC content representation. Enzymatic cytosine conversion has been proposed as an alternative method for methylation mapping. Compared to bisulfite sequencing, enzymatic methylation sequencing (EM-seq) has more balanced genome coverage with less GC-bias and identifies more genomic features with the same number of reads, boosting signal-to-noise ratios in low-input samples.^4^

Compared with whole-genome methylation sequencing, reduced representation methylation sequencing enriches DNA samples for CpG-rich genomic regions via restriction enzyme (e.g., MspI) digestion. This enrichment step prior to cytosine conversion greatly enhances sample throughput and reduces sequencing costs, allowing sample multiplexing and profiling of a larger sample size to enhance statistical power.^5^ A previous study by Vaisvila and colleagues^4^ demonstrated EM-seq’s superior performance in whole-genome methylation sequencing compared with WGBS, yet whether the enzymatic cytosine conversion technology could be adapted for reduced representation methylation sequencing remains unknown.

In this study, we performed methylation profiling of mouse splenic T cell subsets and alveolar T cell subsets from patients with severe SARS-CoV-2 pneumonia to compare the performance of bisulfite-based and enzymatic methylation sequencing methods, including a novel reduced representation EM-seq (RREM-seq) approach. We chose specific T cell subsets—CD4+ conventional T (Tconv) cells and FOXP3+ regulatory T (Treg) cells—that we and others demonstrated to have distinct methylation profiles in mice and humans.^6–9^ Specifically, Treg cells are hypomethylated compared with CD4+ conventional T cells within defined Treg-specific super enhancer (Treg-SE) regions, which are lineage-specifying elements that determine Treg cell identify and function in the lung and other organs.^6^ We found that RREM-seq is a reliable method to perform low-input methylation profiling and demonstrated a similar signal-to-noise ratio compared with RRBS in low-input samples.

## Results

To compare the performance of bisulfite conversion and enzymatic conversion in 5mC detection, we performed whole-genome (WGBS and WGEM-seq) and reduced representation (RRBS and RREM-seq) methylation sequencing, leveraging mouse CD4+ Tconv and Treg cells as cell types with known differences in DNA methylation patterns at defined genomic elements. We used our previously published flow cytometry methods^8,9^ to sort live Tconv and Treg cells from the spleens of *Foxp3*^*YFP-Cre*^ mice, in which yellow fluorescent protein (YFP) marks the Treg cell lineage (**Figure 1A**). We defined standard DNA input as >10 ng, low DNA input as 2-10 ng, and ultra-low DNA input as <2 ng. At standard input, all four methods generated DNA methylation libraries with comparable final concentrations (**Figure 1B**). Unmethylated λ-bacteriophage DNA was included in all samples as an internal negative control, with an average assayed CpG methylation rate of 0.35%, indicating an approximately 100% unmethylated cytosine conversion rate in all libraries (**Figure 1C**). At low- and ultra-low-input levels, RREM-seq successfully generated acceptable quality libraries without the need for additional PCR amplification cycles (**Supplemental Figure 1A**). These libraries fell within the expected size range, and primer contamination was not observed. While RRBS failed to generate reliable DNA libraries when using <2-ng inputs, the concentration and quality of the ultra-low-input RREM-seq libraries was comparable to those of RRBS libraries with more than 10-fold higher DNA input (**Supplemental Figure 1B**).

**Figure 1.**
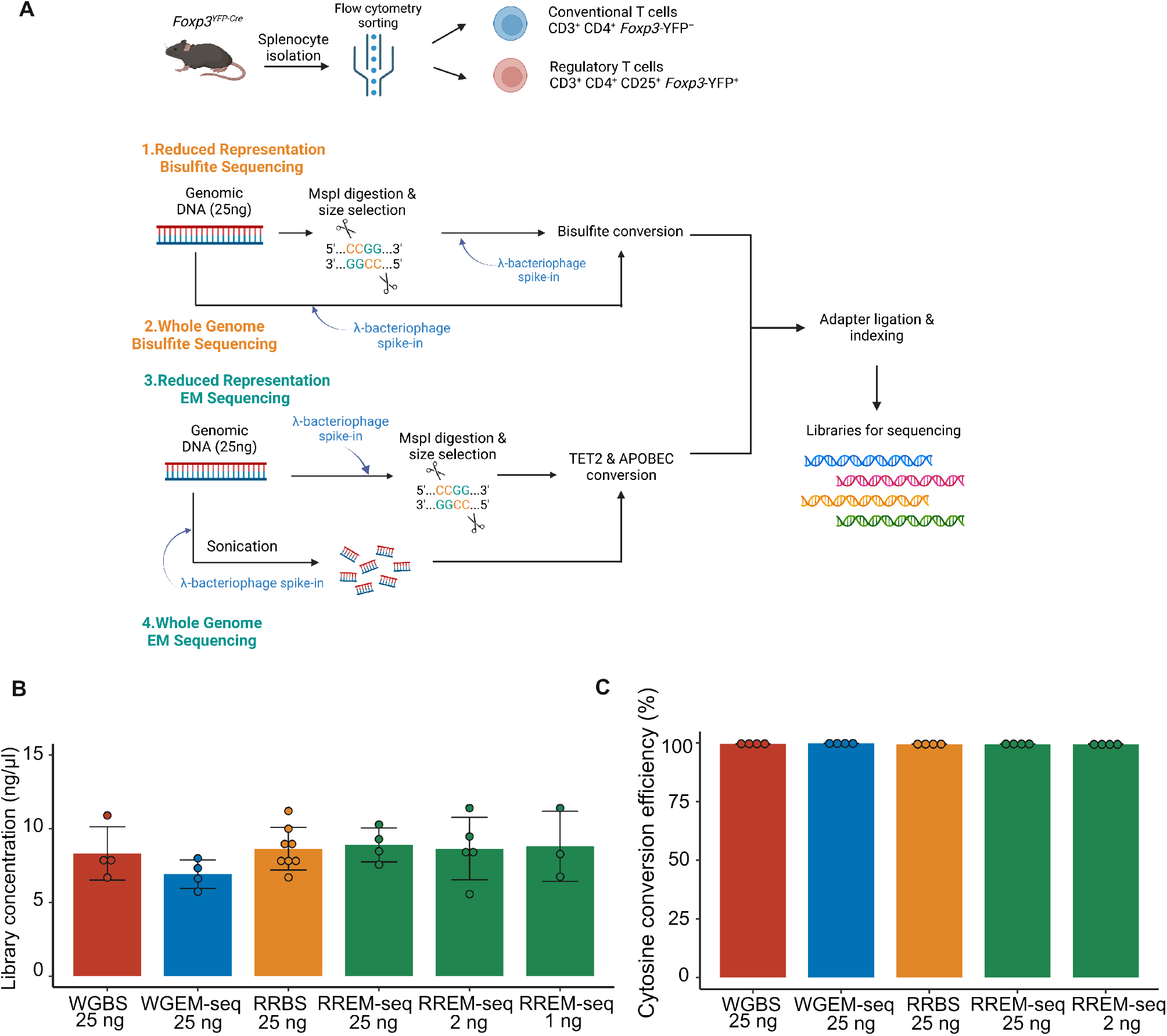
Construction and quality control of DNA methylation libraries. (**A**) Schematic representation of mouse splenic T cell sorting and DNA methylation library construction. (**B-C**) Final library concentration (B) and cytosine conversion efficiency (C) between four different methylation sequencing methods. Error bars show standard deviation.

**Supplemental Figure 1.**
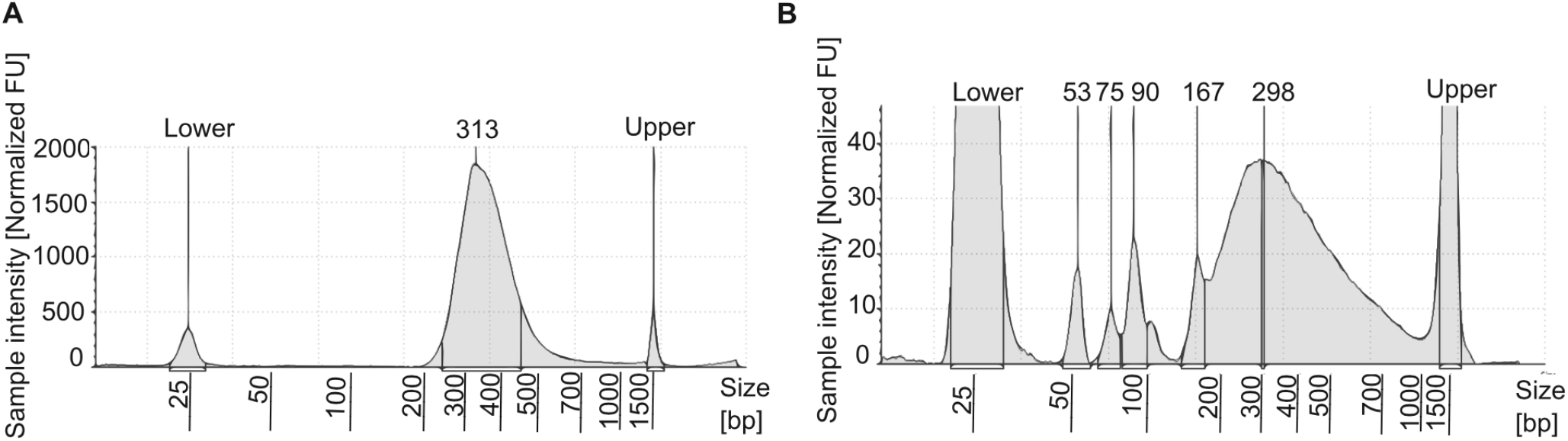
TapeStation traces of low-input DNA methylation libraries. (**A-B**) Final library quality of a representative RREM-seq library generated using 1-ng input (A) and a representative RRBS library using 2-ng input (B). bp, base pair; FU, fluorescence units.

We also compared coverage of genomic elements, including CpG islands (CpGIs) and gene promoters, between the four methylation sequencing methods. At standard input, all four methods (WGBS, WGEM-seq, RRBS, RREM-seq) demonstrated similar coverage of CpGIs and promoters, defined by detection of at least one CpG in the element (**Figure 2A-B**). As inputs decreased from 25 ng to 2 ng, RREM-seq did not demonstrate a decrease in coverage of genomic features, suggesting that the performance of RREM-seq in genomic element detection is not correlated with input concentration at these input levels. We observed average hypomethylation across CpGIs in all sequencing methods (**Figure 2C-F**). Raw quantification of CpG methylation surrounding the transcriptional start site (TSS) also demonstrated the classic dip-and-plateau trend,^10^ indicating hypomethylation around the TSSs and hypermethylation within gene bodies (**Figure 2G-J**).

**Figure 2.**
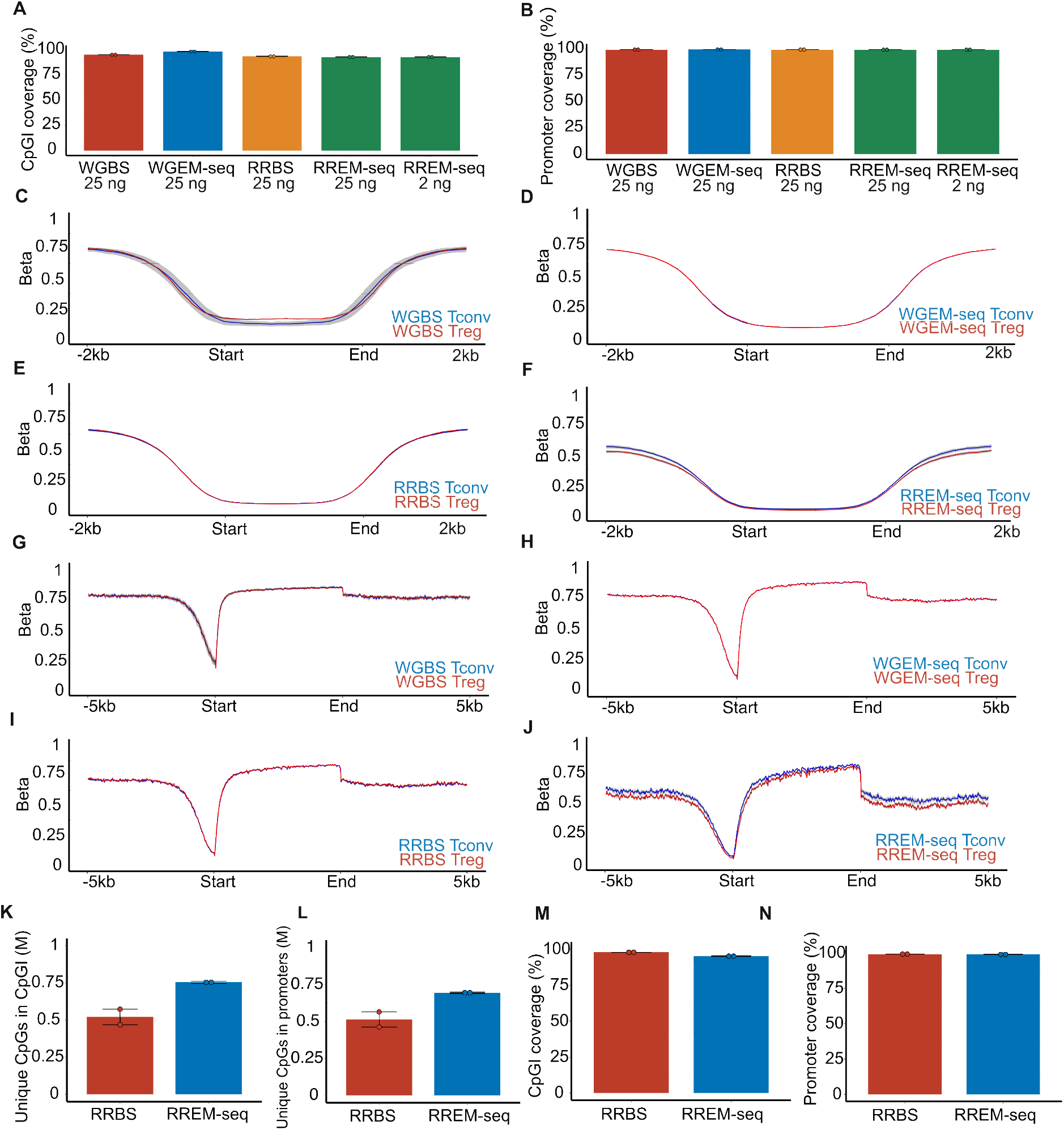
CpG methylation in regulatory genomic elements. (**A-B**) Coverage of CpGIs (A) and promoters (B). (**C-F**) Quantification trend plots of β scores across CpGIs with 2 kb of flanking sequence. (**G-J**) Quantification trend plots of β scores across gene bodies defined by transcriptional start site (TSS) and transcriptional end site (TES) with 5 kb of flanking sequence. Data represent merged average of 2 technical replicates for each cell type, and the shaded area represents the range. (**K-L**) Number of unique CpGs detected in CpGIs (K) and promoters (L) and coverage of CpGIs (M) and promoters (N) using 10% of the original reads in each reduced representation sample. Error bars represent range.

To further assess the performance of RREM-seq in profiling low-read samples, we performed *in silico* random downsampling of the RRBS and RREM-seq libraries. 10% of the total reads from each sample were randomly selected, and we repeated the unique CpG detection and genomic feature coverage analysis in these low read count samples. While the total number of detected CpGIs and promoters were similar between RRBS and RREM-seq (**Figure 2K-L**), RREM-seq was able to detect more unique CpGs in these regulatory genomic elements compared with RRBS (**Figure 2M-N**).

To validate the performance of RREM-seq in identifying methylation differences between Tconv and Treg cells, we quantified methylation levels within the Treg-SE regions.^7^ In both standard-input and low-input RREM-seq libraries, CpG methylation across Treg-SEs demonstrated relative hypomethylation in Treg cells. This trend was not observed in the surrounding regions, confirming RREM-seq’s performance in profiling low-input samples (**Figure 3A-J** and **Supplemental Figure 2**).

**Figure 3.**
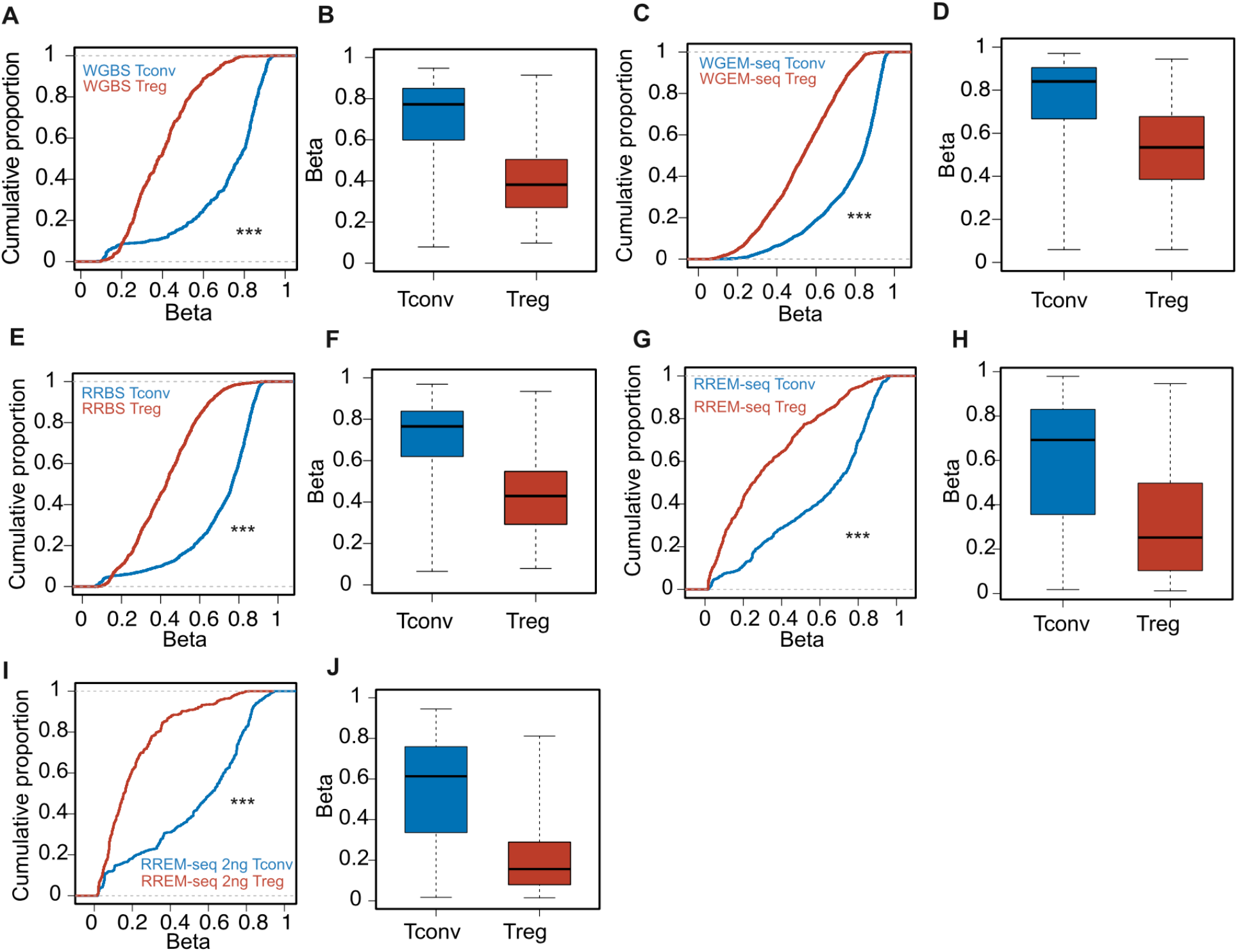
CpG methylation in Treg super-enhancer elements. (**A-J**) β values of differentially methylated CpGs across Treg-SE in Tconv and Treg cells in WGBS (A-B; *n* = 2293), WGEM-seq (C-D; *n* = 8748), RRBS (E-F; *n* = 4476), and RREM-seq (standard input: G-H; *n* = 3245; low input: I-J; *n* = 1579) libraries, FDR *q* < 0.05. In (A, C, E, G, I), the cumulative distribution function of β values across the Treg-SE is plotted against their cumulative proportion. A leftward shift indicates hypomethylation. (B, D, F, H, J) Box-and-whisker plots comparing β values across the Treg-SE between Tconv and Treg cells. Data represent merged average of 2 technical replicates. β values continuously range from 0 (unmethylated) to 1 (methylated). \*\*\**p* < 0.0001 by Kolmogorov–Smirnov test.

**Supplemental Figure 2.**
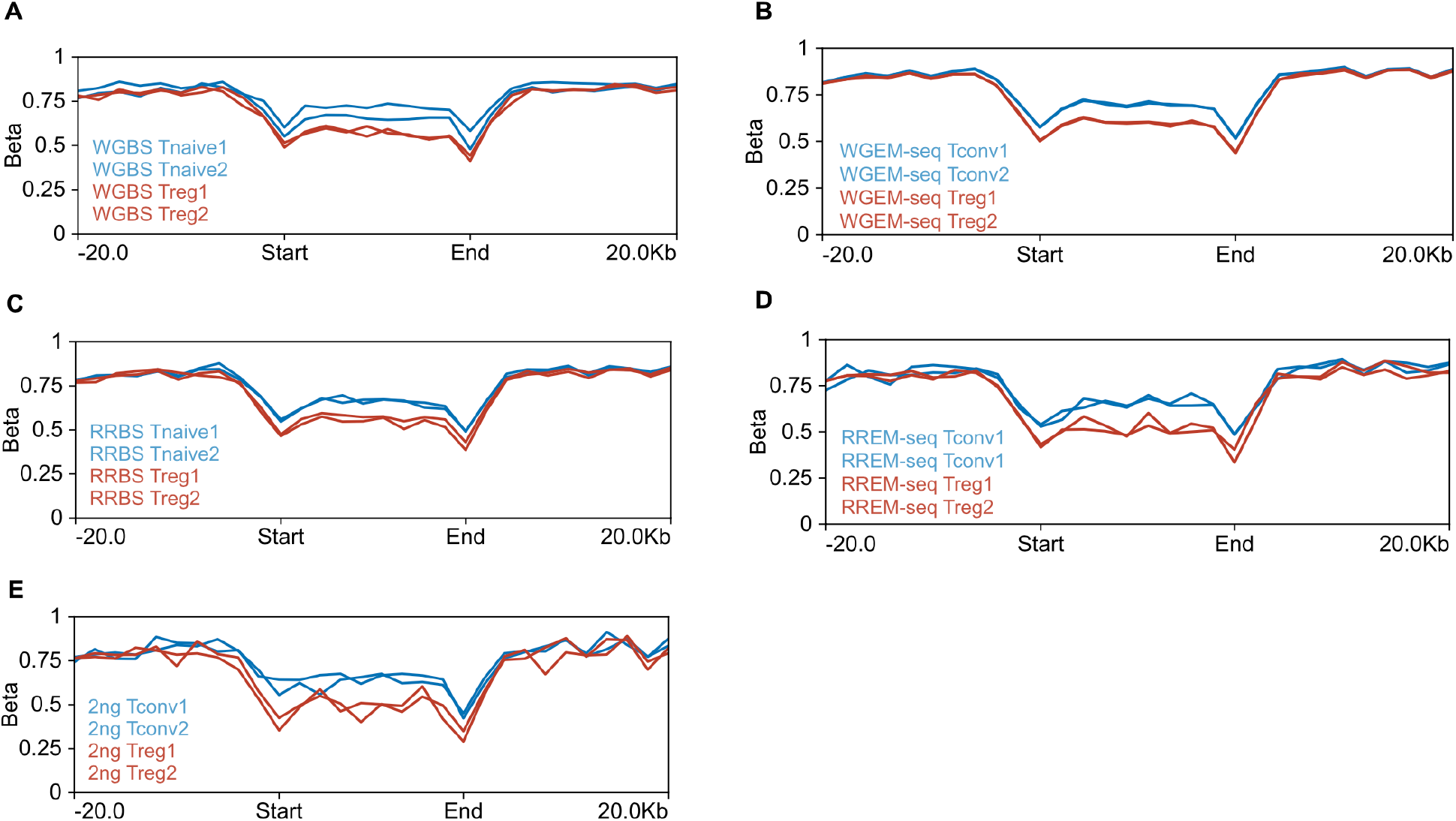
Analysis of Treg super-enhancer elements and surrounding regions. (**A-E**) Quantification trend plot of raw DNA methylation (β scores) of WGBS (A), WGEM-seq (B), RRBS (C), RREM-seq (standard input: D; low input: E) libraries across Treg-SE with 20 kb of flanking sequence on either side.

The T cell response to lower respiratory tract infection is a critical determinant of outcomes in patients with severe pneumonia.^11,12^ To further demonstrate the performance of RREM-seq in assessing low-input clinical samples, we performed methylation profiling of alveolar T cells obtained by bronchoalveolar lavage (BAL) from mechanically ventilated patients with severe SARS-CoV-2 pneumonia within 48 hours of intubation. Alveolar T cells were sorted from BAL fluid samples that were collected as part of the Successful Clinical Response in Pneumonia Therapy (SCRIPT) study, an observational cohort study of mechanically ventilated patients with severe pneumonia.^13^ As described in previous publications,^6,13^ we used flow cytometry to isolate alveolar CD4+ CD127low CD25+ Treg cells and non-Treg CD4+ cells. We obtained 7 paired Tconv (average cell count = 33,584) and Treg (average cell count = 2946) cell samples. Methylation libraries were generated using ≤2 ng genomic DNA (2 ng for Tconv cells and 0.9–2 ng for Treg cells based on sample availability) via our optimized RREM-seq procedures. On average, the final library concentration was 6.8 ng/μL (**Figure 4A**), which is similar to the libraries generated using standard input mouse splenic T cells above. Average CpG methylation was 0.5% for unmethylated λ-bacteriophage DNA, indicating an approximately 100% unmethylated cytosine conversion rate (**Figure 4B**). We observed no significant correlation between sample DNA input and final library concentration (Pearson correlation coefficient = -0.12, *p* = 0.68) (**Figure 4C**). These RREM-seq libraries also demonstrated similar coverage of CpGIs and promoters to standard-input RRBS libraries, confirming RREM-seq’s performance in profiling low-input clinical samples (**Figure 4D-G**).

**Figure 4.**
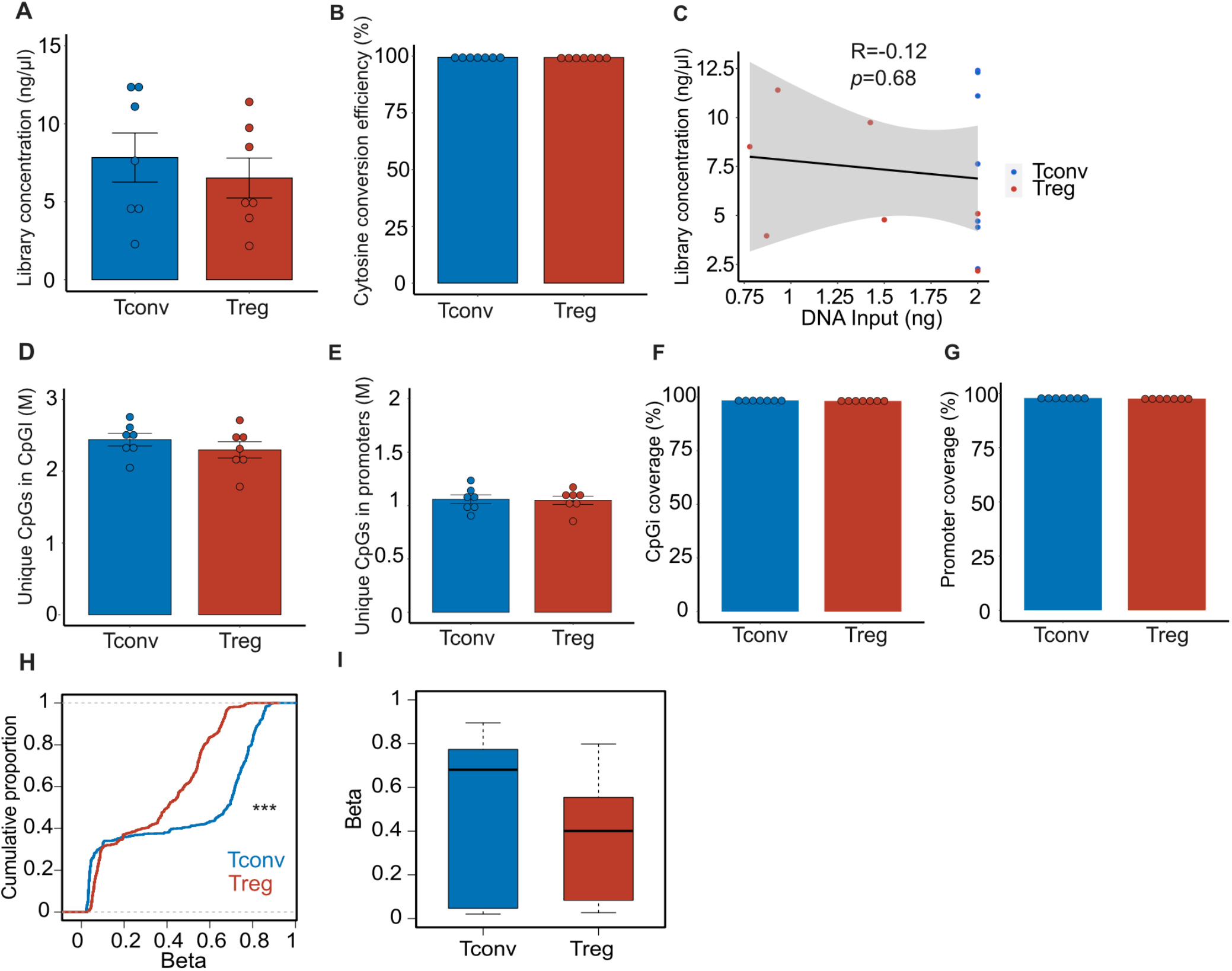
Methylation profiling of ultra-low-input alveolar T cell samples from patients with severe SARS-CoV-2 pneumonia. (**A-B**) Final library concentration (A) and cytosine conversion efficiency (B) in alveolar Tconv and Treg cells. (**C**) Correlation plot comparing RREM-seq library concentration with sample DNA input. The black line and shaded area represent the linear regression fit line with 95% confidence interval. (**D-E**) Unique CpGs detected in CpGIs (D) and promoters (E). (**F-G**) Coverage of CpGIs (F) and promoters (E). (**H**) Cumulative distribution function comparing β values of differentially methylated CpGs (*n* = 1759) within Treg-SE between alveolar Tconv and Treg cells, FDR *q* < 0.05. (**I**) β values of differentially methylated CpGs within Treg-SE. Data represent merged average of 7 replicates for each cell type obtained from 7 individual patients. \*\*\**p* < 0.0001 by Kolmogorov–Smirnov test.

**Supplemental Figure 3.**
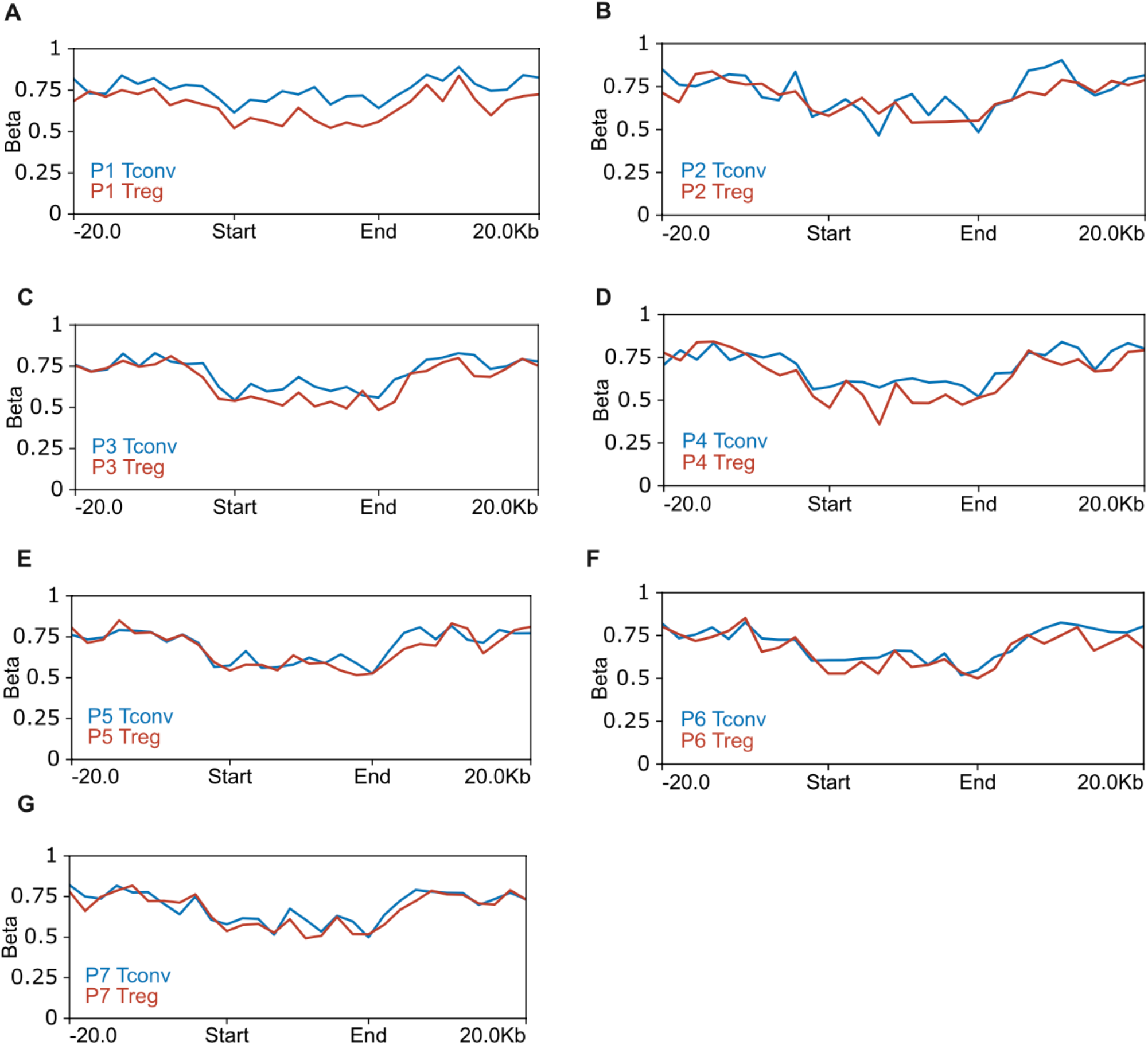
Analysis of Treg super-enhancer elements in patients with severe SARS-CoV-2 pneumonia. (**A-G**) Quantification trend plot of raw DNA methylation (β scores) across Treg-SE with 20 kb of flanking sequence on either side from each patient.

We previously demonstrated that the distinct Treg-SE DNA methylation patterns present in mouse Treg and Tconv cells are also found in Treg and Tconv cells in BAL fluid obtained from patients with severe pneumonia.^6^ To compare methylation status within the Treg-SE regions in alveolar Tconv and Treg cells, we mapped 62 out of 66 regulatory elements from the original mm9 mouse reference genome onto the hg38 human reference genome. Alveolar Treg cells demonstrated hypomethylation across the Treg-SE regions compared with Tconv cells (**Figure 4H-I**). These results in ultra-low-input samples are consistent with our previous publication,^6^ in which we used RRBS to compare methylation levels between alveolar Tconv cells and Treg cells across the Treg-SE regions in standard input samples. We also compared raw quantification of CpG methylation of Tconv and Treg cells in each patient; the Treg-SE DNA methylation pattern varied between patients (**Supplemental Figure 3**).

## Discussion

Due to fewer sequencing reads required per sample, reduced representation methods offer a cost-effective alternative to whole-genome methylation sequencing approaches.^3^ While RRBS and RREM-seq show comparable performance at standard input levels, our results demonstrate that RREM-seq outperforms RRBS at ultra-low input levels at which RRBS could not generate high quality libraries. We found that when using <2 ng of DNA, RREM-seq generates reliable DNA methylation libraries and has comparable performance against standard-input RRBS libraries in detecting CpGs across regulatory genomic elements and gene bodies. We also demonstrated RREM-seq’s performance in profiling methylation status in ultra-low-input clinical samples, determining that RREM-seq detects Treg cell-specific hypomethylation patterns in alveolar Treg cells compared with alveolar Tconv cells in patients with severe SARS-CoV-2 pneumonia.

Bisulfite sequencing, the current gold standard, is limited by several important caveats. Bisulfite treatment involves deamination of cytosine residues in single-stranded DNA at 37–50°C followed by bisulfite removal using extreme pH adjustment (pH > 13).^14^ These chemical reactions result in substantial DNA degradation via depyrimidination.^14^ On top of this bisulfite-induced DNA degradation, the CpG enrichment step in reduced representation protocols has a yield of around 5–10% of the original input after DNA fragment size selection, further lowering DNA input. Hence, the required DNA input for RRBS may not be feasible for many clinical samples and rare cell populations. Compared to bisulfite treatment, enzymatic conversion of cytosine residues does not contribute to DNA degradation, the conversion process does not require extreme pH, and the reaction temperature is kept at no higher than 37°C.^4^ In addition to 5mC detection, the EM-seq pipeline can be modified to specifically identify 5hmC by incorporating the enzyme T4-GBT for 5hmC glycosylation.

Similar to the bisulfite-based methods, the accuracy of CpG detection heavily relies on the efficient conversion of unmethylated cytosine residues to uracil residues. Unconverted cytosine residues are categorized as 5mC by the downstream analysis and therefore result in inaccurate methylation profiling. Our results confirmed that enzymatic conversion is highly efficient in converting unmethylated cytosines. Regardless, unmethylated negative control DNA (for example λ-bacteriophage) should be incorporated into the RREM-seq pipeline to address this potential limitation.

Overall, our results demonstrate that RREM-seq is a reliable method to perform methylation profiling and displays superior signal-to-noise ratios compared with RRBS in ultra-low-input samples. Many low-input samples have great potential to answer relevant research questions, yet the current RRBS pipeline cannot generate reliable methylation libraries using ultra-low input levels. Combining a reduced representation strategy with enzymatic conversion technology provides an opportunity to perform cost-effective methylation profiling of biologically relevant yet limited cell populations, especially in clinical samples.

## Methods

### Mouse tissue preparation and flow cytometry sorting

Spleens were harvested from healthy *Foxp3*^*YFP-Cre*^ mice (Jackson Labs Strain #:016959), in which yellow fluorescent protein (YFP) marks the Treg cell lineage. Single-cell suspensions were then prepared, and red blood cells were removed via ACK Lysis Buffer (ThermoFisher). Staining was performed using reagents listed in **Supplemental Table 1**. Cell sorting of conventional T (CD3ε+ CD4+ *Foxp3*-YFP–) and Treg (CD3ε+ CD4+ CD25+ *Foxp3*-YFP+) cells was performed using BD Biosciences FACSAria SORP instruments with FACSDiva software.

**Supplemental Table 1.** Flow cytometry fluorochromes used for sorting mouse T cells.

| Reagent/Antigen | Conjugate | Clone | Company | Catalog no. |
| --- | --- | --- | --- | --- |
| CD3ε | PE | 145-2C11 | eBioscience | 12-0031-83 |
| CD4 | PerCP-Cy5.5 | GK1.5 | Biolegend | 100434 |
| CD25 | APC | PC61.5 | Invitrogen | 2154040 |
| Fixable Viability Dye eFluor 506 | N/A | N/A | eBioscience | 65-0866-14 |

### T cell samples from patients with severe SARS-CoV-2 pneumonia

Bronchoalveolar lavage (BAL) samples were collected from mechanically ventilated patients with severe SARS-CoV-2 pneumonia within 48 hours of intubation. All patients were enrolled in the Successful Clinical Response In Pneumonia Therapy (SCRIPT) study under Northwestern University IRB STU00204868. For SCRIPT participants, the first BAL (study day 0) is performed within 48 hours following intubation. The etiology of pneumonia was determined based on clinical and BAL fluid analysis by consensus of five pulmonary and critical care medicine physicians.^15^ All BAL samples were sorted and analyzed using our published flow cytometry methods^6^ and then stored at -80°C. In this study, we included the sorted day 0 alveolar Treg cells (live FSClo SSClo CD3ε+ CD4+ CD25hi CD127lo) and Tconv cells (live FSClo SSClo CD3ε+ CD4+ cells not in the Treg cell gate)^6^ from the SCRIPT study and performed methylation profiling using RREM-seq.

### Methylation library generation

Flow cytometry-sorted cells were first lysed with QIAGEN RLT Plus and then genomic DNA was extracted using the AllPrep DNA/RNA Micro Kit (QIAGEN). For WGEM-seq libraries, 25 ng of genomic DNA was first fragmented to a size of 240–290-bp using a Covaris E220 sonicator and then enzymatically converted with TET2 and APOBEC (New England Biolabs) per the manufacturer’s instructions. For WGBS libraries, 25 ng of genomic DNA was bisulfite converted with the EZ DNA Methylation-Lightning Kit (Zymo Research) per the manufacturer’s protocol. An additional restriction enzyme digestion step was included for RRBS and RREM-seq libraries, in which genomic DNA was fragmented with MspI (New England Biolabs) and then size-selected for 100–250-bp fragments using solid-phase reversible immobilization beads (MagBio Genomics).^16^ CpG-enriched genomic DNA was then bisulfite converted for RRBS libraries and enzymatically converted for RREM-seq libraries.

Random priming, adapter ligation, PCR product clean-up, and final library amplification were performed using the NEBNext Enzymatic Methyl-seq Kit (New England BioLabs) for WGEM-seq libraries, and the Pico Methyl-Seq Library Prep Kit (Zymo Research) was used for WGBS, RRBS, and RREM-seq libraries. Bisulfite-based methods (WGBS and RRBS) and enzyme-based methods (WGEM-seq and RREM-seq) required 10 and 8 PCR cycles for final amplification, respectively. Final library size distribution and quality were assessed via high-sensitivity screen tape (TapeStation 4200, Agilent). Unmethylated λ-bacteriophage DNA (1:200 mass ratio; New England Biolabs) was included in all samples to calculate unmethylated cytosine conversion efficiency. Four-to-six libraries were pooled in an equimolar ratio prior to sequencing. Whole-genome libraries were sequenced using paired-end reads with a NextSeq 2000 P3 reagent kit (200 Cycles; Illumina) and reduced representation libraries were sequenced using single-end reads with a NextSeq 2000 P2 reagent kit (100 Cycles; Illumina).

### Methylation library analysis

Methylation sequencing analysis was performed using our published Bismark-based pipelines.^5,6,9^ After sequencing, raw binary base call (BCL) files were converted to FASTQ files using bcl-Convert (version 3.10.5; Illumina) and then trimming was performed using Trim Galore! (version 0.4.3). Bismark (version 0.21.0) was used to perform alignment to the reference genomes (mm10 or hg38 assembly according to the sample source) and methylation extraction. Bismark coverage files were used for quantification using SeqMonk (version 1.48.0) and R package DSS (version 2.46.0). For genomic feature coverage analysis, CpG islands, transcription start site, and transcription end site coordinates were obtained from the UCSC mm10/hg38 annotation tracks. CpG shores were defined as 2-kb regions flanking CpG islands and promoters were defined as 1-kb regions flanking transcription start sites. *In silico* downsampling of FASTQ files was performed using the seqtk toolkit to randomly sample 10% of the total reads after adapter trimming and QC filtering (samtools view -s 0.1 -b).

### Treg-SE

Published coordinates of Treg-SE elements^7^ were first mapped from the original mm9 to the mm10 reference genome and then mapped to the hg38 reference genome using the UCSC Batch Coordinate Conversion (liftOver) tool with the default settings (minimum ratio of bases that must remap = 0.1).

### Statistics

All statistical analyses were performed in R (version 4.2.3) and SeqMonk (version 1.48.0). All computational processing and analysis were performed using Northwestern University’s High-Performance Computing Cluster. A *p* value or false discovery rate (FDR) *q* value < 0.05 was considered statistically significant.

## Data and code availability

Raw sequencing data of the mouse samples and all processed CpG coverage files will be made publicly available via the GEO repository pending peer-reviewed publication. Raw sequencing data of the patient samples will be made available via dbGaP. Code used for differential methylation analysis and Treg-SE analysis is available at https://github.com/palupaca/RREMseq_2024.

## Acknowledgements

QL is supported by the David W. Cugell Fellowship and the Genomics Network (GeNe) Pilot Project Funding. CPRF is supported by T32HL076139. NM is supported by NIH award T32AI083216. AMJ is supported by NIH award F32HL162418. LM-N is supported by the NIH awards K08HL159356, U19AI135964, and the Parker B. Francis Opportunity Award. BDS is supported by NIH awards R01HL149883, R01HL153122, P01HL154998, P01AG049665, U19AI135964, and U19AI181102. We wish to acknowledge the Northwestern University Flow Cytometry Core Facility supported by CA060553; the BD FACSAria SORP system was purchased with the support of S10OD011996. We also wish to acknowledge the Northwestern University Metabolomics and Integrative Genomics Core. This research was supported in part through the computational resources and staff contributions provided by the Genomics Compute Cluster, which is jointly supported by the Feinberg School of Medicine, the Center for Genetic Medicine, and Feinberg’s Department of Biochemistry and Molecular Genetics, the Office of the Provost, the Office for Research, and Northwestern Information Technology. The Genomics Compute Cluster is part of Quest, Northwestern University’s high performance computing facility, with the purpose to advance research in genomics. Figure 1A was created using biorender.com.

## References

1. Schübeler, D. Function and information content of DNA methylation. Nature 517, 321–326 (2015).

2. Saxonov, S., Berg, P. & Brutlag, D. L. A genome-wide analysis of CpG dinucleotides in the human genome distinguishes two distinct classes of promoters. Proc. Natl. Acad. Sci. 103, 1412–1417 (2006).

3. Meissner, A. et al. Reduced representation bisulfite sequencing for comparative high-resolution DNA methylation analysis. Nucleic Acids Res. 33, 5868–5877 (2005).

4. Vaisvila, R. et al. Enzymatic methyl sequencing detects DNA methylation at single-base resolution from picograms of DNA. Genome Res. 31, 1280–1289 (2021).

5. Singer, B. D. A Practical Guide to the Measurement and Analysis of DNA Methylation. Am. J. Respir. Cell Mol. Biol. 61, 417–428 (2019).

6. Walter, J. M., Helmin, K. A., Abdala-Valencia, H., Wunderink, R. G. & Singer, B. D. Multidimensional assessment of alveolar T cells in critically ill patients. JCI Insight 3, (2018).

7. Kitagawa, Y. et al. Guidance of regulatory T cell development by Satb1-dependent super-enhancer establishment. Nat. Immunol. 18, 173–183 (2017).

8. Helmin, K. A. et al. Maintenance DNA methylation is essential for regulatory T cell development and stability of suppressive function. J. Clin. Invest. 130, 6571–6587.

9. Morales-Nebreda, L. et al. Aging imparts cell-autonomous dysfunction to regulatory T cells during recovery from influenza pneumonia. JCI Insight 6, (2021).

10. Jjingo, D., Conley, A. B., Yi, S. V., Lunyak, V. V. & Jordan, I. K. On the presence and role of human gene-body DNA methylation. Oncotarget 3, 462–474 (2012).

11. Markov, N. S. et al. A distinctive evolution of alveolar T cell responses is associated with clinical outcomes in unvaccinated patients with SARS-CoV-2 pneumonia. 2023.12.13.571479 Preprint at 10.1101/2023.12.13.571479 (2023).

12. Singer, B. D. & Chandel, N. S. Immunometabolism of pro-repair cells. J. Clin. Invest. 129, 2597–2607 (2019).

13. Grant, R. A. et al. Circuits between infected macrophages and T cells in SARS-CoV-2 pneumonia. Nature 590, 635–641 (2021).

14. Tanaka, K. & Okamoto, A. Degradation of DNA by bisulfite treatment. Bioorg. Med. Chem. Lett. 17, 1912–1915 (2007).

15. Pickens, C. I. et al. An Adjudication Protocol for Severe Pneumonia. Open Forum Infect. Dis. 10, ofad336 (2023).

16. McGrath-Morrow, S. A. et al. DNA methylation regulates the neonatal CD4+ T-cell response to pneumonia in mice. J. Biol. Chem. 293, 11772–11783 (2018).

